# Structural basis of Q-dependent antitermination

**DOI:** 10.1101/651190

**Authors:** Zhou Yin, Jason Kaelber, Richard H. Ebright

## Abstract

Lambdoid bacteriophage Q protein mediates the switch from middle to late bacteriophage gene expression by enabling RNA polymerase (RNAP) to read through transcription terminators preceding bacteriophage late genes. Q loads onto RNAP engaged in promoter-proximal pausing at a Q binding element (QBE) and an adjacent sigma-dependent pause element (SDPE) to yield a “Q-loading complex,” and Q subsequently translocates with RNAP as a pausing-deficient, termination-deficient “Q-loaded complex.” Here, we report high-resolution structures of four states on the pathway of antitermination by Q from bacteriophage 21 (Q21): Q21, the Q21-QBE complex, the Q21-loading complex, and the Q21-loaded complex. The results show that Q21 forms a torus--a “nozzle”--that narrows and extends the RNAP RNA-exit channel, extruding single-stranded RNA and preventing formation of pause and terminator hairpins.

**One Sentence Summary:** Q forms a “nozzle” that narrows the RNA polymerase RNA-exit channel and extrudes ssRNA, preventing formation of RNA hairpins.

## Main Text

Lambdoid bacteriophage Q protein regulates gene expression through transcription antitermination, a mechanism of regulation that is widely used in bacteria and bacteriophage but that has been poorly understood (1–10; reviewed in 11-13).

Q mediates the temporal switch from middle to late bacteriophage gene expression by enabling RNA polymerase (RNAP) to read through a transcription terminator preceding bacteriophage late genes (Fig. 1A; 1–13). The Q-dependent gene regulatory cassette consists of the gene for Q followed by a transcription unit comprising a promoter (PR’), a promoter-proximal σ-dependent pause element (SDPE; a sequence resembling a promoter −10 element, at which σR2, the σ-factor module that recognizes promoter −10 elements, re-establishes sequence-specific protein-DNA interactions), and a terminator, followed by bacteriophage late genes. In the absence of Q, RNAP initiating transcription at the PR’ promoter pauses at the SDPE and terminates at the terminator, and late genes are not expressed; in the presence of Q, RNAP initiating at the PR’ promoter, escapes the SDPE and reads through the terminator, and late genes are expressed.

**Fig. 1.**
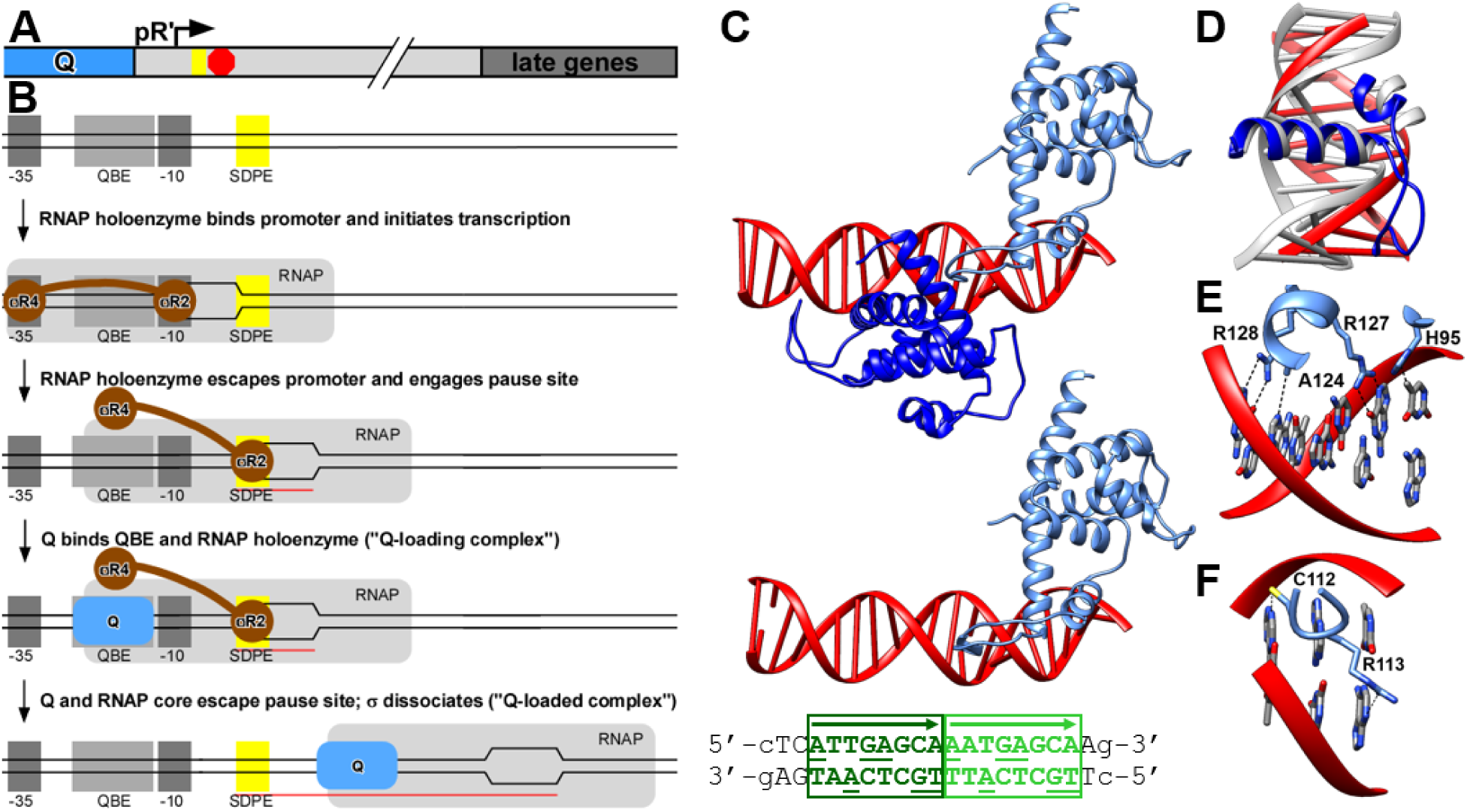
Biological function of Q and structure of Q21-QBE complex. **(A)** Q-dependent regulatory cassette, consisting of gene for Q (blue) and adjacent transcription unit comprising pR’ promoter (arrow), SDPE (yellow rectangle), terminator (red octagon), and bacteriophage late genes (gray). **(B)** Steps in assembly and function of a Q-dependent transcription antitermination complex. Promoter −35 and −10 promoter elements, dark gray rectangles; QBE, light gray rectangle; SDPE, yellow rectangle; RNAP core enzyme, large light grey rectangle; σ, brown; Q, blue; DNA nontemplate and template strands, black lines (unwound transcription bubble indicated by raised and lowered line segments); RNA, red line. **(C)** Crystal structure of Q21-QBE complex. Top, molecular assembly in asymmetric unit comprising two Q protomer interacting with QBE. Middle, molecular assembly in asymmetric unit comprising one Q protomer interacting with QBE. Bottom, QBE DNA fragment (QBEu, dark green; QBEd, light green). **(D)** Q21 HT[loop]T motif interacting with DNA (blue and red, superimposed on HTH motif of ECF σ factor σR4 interacting with DNA (PDB 2H27; gray). **(E)** Interactions of Q21 with DNA major groove (top) and DNA minor groove (bottom).

Q is targeted to the PR’ transcriptional unit through a multi-step process entailing: (i) formation of a “Q-loading complex” comprising a Q protein bound to a Q binding element (QBE) and a σ-containing transcription elongation complex (TEC) paused at the SDPE, and (ii) transformation into a “Q-loaded complex” comprising a Q-containing TEC that translocates processively, ignores pause elements, and ignores terminators (Fig. 1B; 3–13).

Here, we report a set of four structures that define the structural basis of antitermination by Q from lambdoid bacteriophage Q21 (Q21; 11,14-15): i.e., (i) a crystal structure of Q21, (ii) a crystal structure of the Q21-QBE complex, (iii) a cryo-EM structure of the Q-loading complex, and (4) a cryo-EM structure of the Q-loaded complex.

We solved a crystal structure of Q21 at 1.9 Å resolution by use of single-wavelength anomalous dispersion (Fig. S1; Table S1). The structure reveals that the fold of Q21 is related to the fold of the σ-factor module that recognizes promoter −35 elements, σR4, and is particularly related to the fold of σR4 from extracytoplasmic-function (ECF) σ factors (Fig. S1C) and of σR4 from ECF σ factors bound to anti-σ factors (Fig. S1D; DALI Z scores ≥6.0; 16-17). σR4 binds DNA through a helix-turn-helix motif (HTH; red in Fig. S1C-D; 16-17). The structural alignment of Q21 and σR4 indicates that Q21 contains an HTH-related, but novel, helix-turn[loop]-helix motif (HT[loop]H; Fig, S1C-D), suggesting that Q21 binds DNA through an HTH-related, but novel, HT[loop]H.

We solved a crystal structure of the Q21-QBE complex at 2.8 Å resolution by use of molecular replacement (Figs. 1C-E and S2; Table S2). The structure showed two distinct molecular assemblies in the asymmetric unit (Figs 1C and S2A). The first assembly comprised two Q21 protomers (Q-upstream and Q-downstream, Qu and Qd) bound to a direct repeat of two 8-bp subsites (QBE-upstream and QBE-downstream, QBEu and QBEd; Figs. 1C, top and S2A, left; Table S2). The second assembly comprised one Q21 protomer (Qd) bound to one 8-bp subsite (QBEd; Figs. 1C, middle and S2A, right). The observed interactions between Q21 and DNA are identical for Qu-QBEu and Qd-QBEd. In each case, the second α-helix of the Q21 HT[loop]H interacts with the DNA major groove, making direct interactions with DNA base edges (Fig. 1E, top and S2B), and the loop of the Q21 HT[loop]H interacts with the adjacent DNA minor groove, making direct interactions with DNA base edges (Figs. 1E, bottom and S2B). In the assembly comprising two Q21 protomers, Qu and Qd, bound to two subsites, QBEu and QBEd, the loop of the HT[loop]H of Qd makes direct protein-protein interactions with Qu (Fig. S2C). Fluorescence-polarization DNA binding experiments indicate that QBEu alone is a low-affinity subsite, QBEd alone is a moderate-affinity subsite, and the intact QBE, comprising both QBEu and QBEd and enabling cooperative protein-DNA interactions by Q21 protomers binding to adjacent subsites, is a high-affinity site (Fig. S2D).

We determined a single-particle-reconstruction cryo-EM structure of the Q21-loading complex at 3.5 Å resolution (Figs. 2 and S3-S4; Table S2). We prepared the Q21-loading complex by *in vitro* reconstitution from recombinant Q21, recombinant *Escherichia coli* RNAP core, recombinant *E. coli* σ^70^, and a synthetic nucleic-acid scaffold containing the PR’21 QBE, a consensus version of the PR’21 SDPE, and an 11 nt RNA (Fig. 2A). The DNA duplex contained a 13 bp non-complementary region overlapping the SDPE, corresponding to the unwound transcription bubble in a TEC at the point of initial pause capture, prior to subsequent cycles of scrunching and backtracking (18-22). The structure shows two Q21 protomers, Qu and Qd, interacting with QBEu and QBEd in a manner matching that in the crystal structure of Q21-QBE (Fig. 1C, top) and simultaneously interacting with a σ-containing TEC (Figs. 2B and S4A-B). σR2 and σR3 make interactions with RNAP and DNA equivalent to those in a transcription initiation complex. In contrast, σR4 and the σR3-R4 linker are disengaged from their positions in a transcription initiation complex, where they occupy the RNAP dock and the RNAP RNA-exit channel (16), and are disordered (Fig. 2B), consistent with the expectation that interactions of σR4 and the σR3-R4 linker with RNAP are weakened or disrupted upon synthesis of a >10 nt RNA product, entry of the RNA 5’-end into the RNAP RNA-exit channel, and promoter escape (Fig. 1B, line 3; 16). Clear, traceable density is present for the entire 11 nt RNA: the RNA 5’-end nucleotide and one additional RNA nucleotide (positions −11 and −10) are in the RNAP RNA-exit channel, and the other nine RNA nucleotides (position −9 through −1) are base-paired to the transcription-bubble template DNA strand as an RNA-DNA hybrid (Fig. 2B,E-F).

**Fig. 2.**
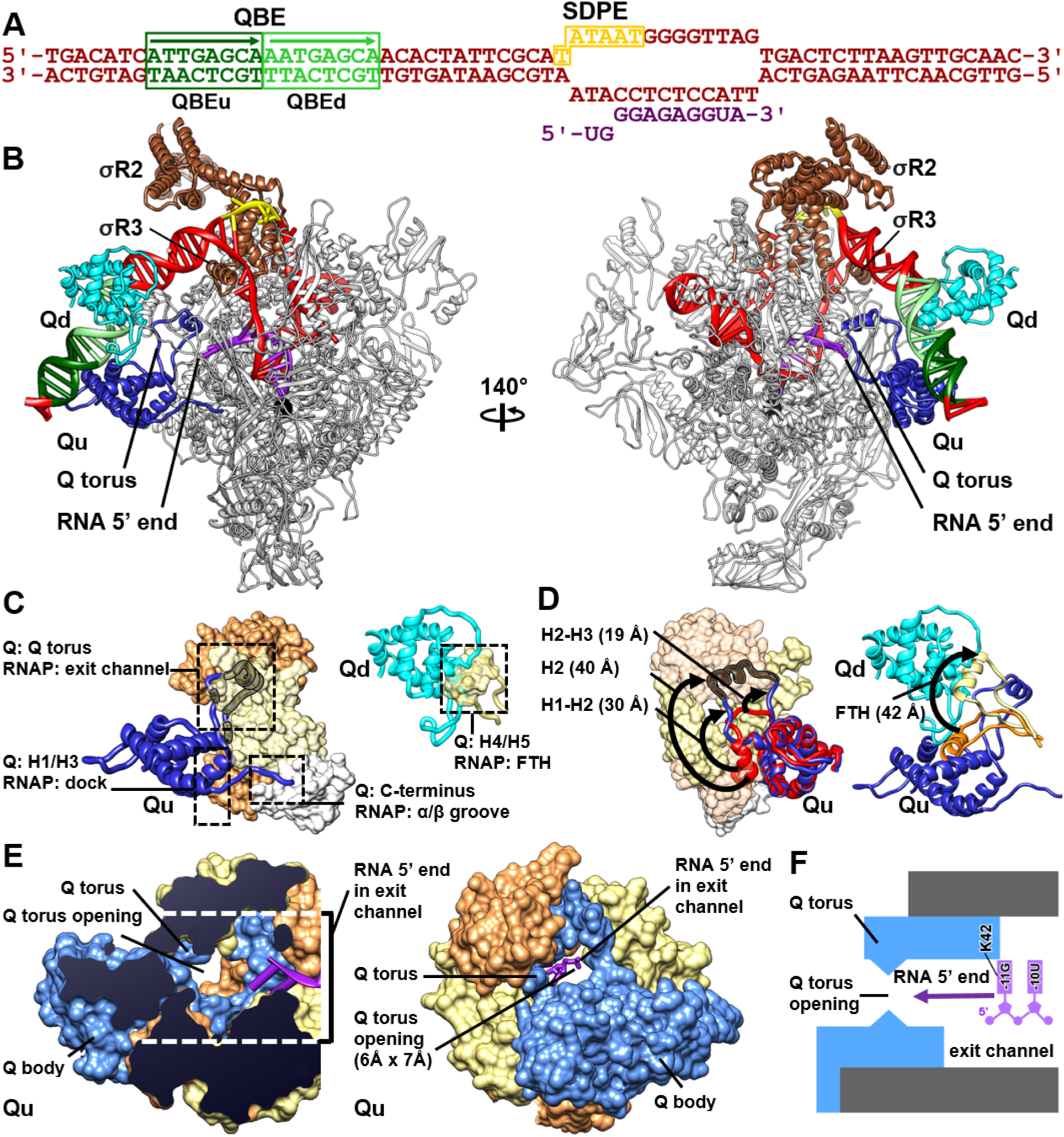
Structure of Q21-loading complex. **(A)** Nucleic-acid scaffold. DNA, red (QBEu, QBEd, and SDPE in dark green, light green, and yellow; non-complementary region corresponding to unwound transcription bubble indicated by raised and lowered letters); RNA, purple. **(B)** Cryo-EM structure of Q21-loading complex (two view orientations). Qu, dark blue; Qd, light blue; RNAP, gray; σ^70^, brown; DNA and RNA, colored as in A; RNAP active-center Mg^2+^, black sphere. **(C)** Qu-RNAP (left) and Qd-RNAP (right) interactions. RNAP β’, β, and α subunits in salmon, tan, and white. Qu segments located inside RNAP RNA-exit channel, behind RNAP surfaces, indicated as gray ribbons with black outlines. Other colors as in B. **(D)** Conformational changes in Qu (left) and RNAP FTH (right) upon formation of Q21-loading complex. Left, Qu in Q-QBE complex (Fig. 1C; red) superimposed on Q21-loading complex (colored as in C). Right; RNAP FTH \in transcription initiation complex (PDB 4YLN; orange) superimposed on Q21-loading complex (colored as in C). **(E)** Restriction of RNAP RNA-exit channel by Q torus and proximity of RNA 5’ end to Q torus (RNA nucleotides numbered with RNA 3’ nucleotide as −1; two view orientations). **(F)** Summary of protein-RNA interactions by Q torus (blue) and RNAP RNA-exit channel (gray).

Both Qu and Qd interact with RNAP (Figs. 2B-C). Qu makes three interactions with RNAP: (i) a “Q torus” comprising the H1-H2 linker, H2, and the H2-H3 linker of Qu interacts at the mouth of, and inside, the RNAP RNA-exit channel; (ii) H3 of Qu interacts with the RNAP dock in manner reminiscent of that for interaction of σR4 with the RNAP dock in a transcription initiation complex (16); and (iii) six C-terminal residues of Qu interact with the RNAP α^I^/β-interface (Fig. 2C, left). The total surface area buried upon Qu-RNAP interaction is 4,657 A^2^, indicative of a high-affinity interaction (23). Qd makes one interaction with RNAP: Qd interacts with the RNAP flap-tip helix (FTH;; Fig. 2C, right). The total surface area buried upon Qd-RNAP interaction is 1,293 A^2^, indicative of a low-affinity interaction (23).

Formation of the Q21-loading complex entails remarkable large-scale conformational changes in Qu and in the RNAP FTH (Fig. 2D). Comparison of the structures of Qu in the Q21-QBE complex and in the Q21-loading complex shows that, upon formation of the Q21-loading complex, the H1-H2 linker, H2, and the H2-H3 linker of Qu move by 19 Å, 40 Å, and 30 Å, respectively, to form the Q torus and to position the Q torus at the mouth of, and inside, the RNAP RNA-exit channel (Fig. 2D, left), and six C-terminal residues of Qu undergo a disorder-order transition to engage the RNAP α^I^/β interface (Fig. 2D, left). Comparison of the structures of the RNAP FTH in a transcription initiation complex and in the Q21-loading complex shows that, upon formation of the Q21-loading complex, the RNAP FTH moves by 42 Å to vacate space to be occupied by Qu and to interact with Qd (Fig. 2D, right)

The binding of the Q torus at the mouth of, and inside, the RNAP RNA-exit channel dramatically restricts the RNAP RNA-exit channel, narrowing and extending the channel (Fig. 2E-F). The Q torus has an opening with a solvent-excluded diameter of just 5-7 Å (Fig. 2G). The structure of the Q21-loading complex indicates that the presence of the Q torus at, and inside, the mouth of the RNAP RNA-exit channel does not affect accommodation of an 11 nt RNA product, and model building indicates it also would not affect accommodation of 12-14 nt RNA products. However, model building indicates that the presence of the Q torus at, and inside, the mouth of the RNAP RNA-exit channel would necessitate >14 nt RNA products: (i) to thread through the Q torus opening (threading RNA nucleotides, with solvent-excluded diameter of 5-7 Å, through the Q-torus opening, with solvent-excluded diameter of 5-7 Å), or (ii), to induce a major change in the conformation or interactions of the Q torus.

We determined a single-particle reconstruction cryo-EM structure of the Q21-loaded complex at 3.8 Å (Figs. 3 and S5-S6; Table S2). To prepare a Q21-loaded complex, we performed *in vitro* transcription with a DNA template comprising the positions −40 to −1 of PR’21 followed by a 70-bp C-less cassette” containing a SDPE at the position of the PR’21 SDPE, followed by a CC-halt site (Fig. 3A), using *E. coli* RNAP σ^70^ holoenzyme, *E. coli* GreB (required for efficient SDPE clearance in the presence of Q21; 11,24), and Q21, in the presence of an NTP subset lacking CTP. 3D-classification of particles showed three major classes of molecular assemblies: (i) Q21-loaded complexes (identifiable by presence of Q21 and absence of σ; 23 %), (ii) TECs (identifiable by absence of Q21 and absence of σ; 21%), (iii) transcription initiation complexes (identifiable by absence of Q21 and presence of σ; 50%) (Fig. S5C). The structure of the class comprising Q21-loaded complexes reveals one Q21 protomer, corresponding to Qu, interacting with a σ-free TEC and having the RNA 5’ end threaded into and through the Q torus (Figs. 3B and S6). The conformation and interactions of Q in the loaded complex are similar to those of Qu in the loading complex (Figs, 2B, 3B). In particular, the Q torus adopts a similar conformation and makes similar interactions with the RNAP RNA-exit channel,, narrowing and extending the RNA-exit channel, in the loaded complex (Figs. 2B, 3B). [The sole notable change is an order-to-disorder transition in the loop of the HT[loop]H motif of Q, associated with the loss of Q-QBE interactions in the loaded complex (Figs. 2B, 3B).]) The structure shows no density for any part of σ (Fig. S6A-B), consistent with the expectation that σ is released upon formation of a loaded complex (Fig. 1B, line 5). Clear, traceable density is present for the 16 nt nucleotides at the 3’ end of the 70 nt RNA product: 2 nt of RNA upstream of the Q torus and outside the RNAP-RNA exit channel (positions −16 and −15), 5 nt of RNA downstream of the Q torus and in the RNAP RNA-exit channel (positions −14 through −10), and 9 nt of RNA base-paired with the transcription-bubble template DNA strand as an RNA-DNA hybrid (positions −9 through −1) (Figs. 2B-D and S6D). An additional unassigned density feature upstream of the Q torus potentially is attributable to additional nucleotides of the 70 nt RNA product (probably positions −21 through −17). The structure thus establishes that, upon formation of the loaded complex, RNA threads into and through the Q torus. The threading of RNA through the Q torus results, in a topological linkage between RNA and Q, creating an effectively unbreakable linkage between the TEC and the Q torus that accounts for the ability of Q to remain stably bound to a TEC (9), resulting in processive antipausing and antitermination over tens of thousands of nucleotide-addition steps (11-13).

**Fig. 3.**
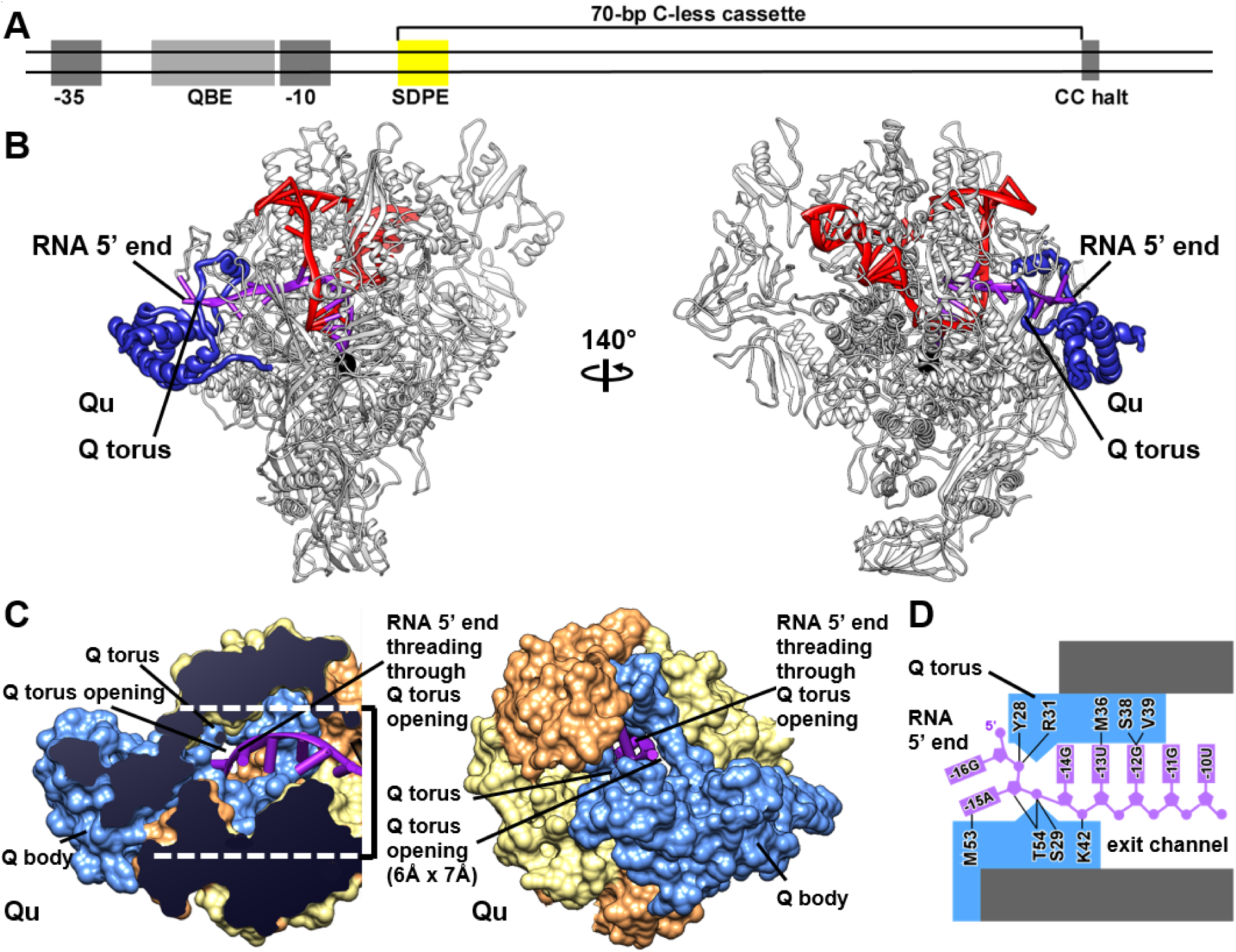
Structure of Q21-loaded complex. **(A)** DNA template for preparation of Q21-loading complex. **(B)** Structure of Q21-loaded complex. View orientations and colors as in Fig. 2B. **(C)** Threading of RNA 5’ end into and through Q torus (RNA nucleotides numbered with RNA 3’ nucleotide as −1). View orientations and colors as in Fig. 2G.. **(D)** Summary of protein-RNA interactions by Q torus (blue) and RNA-exit channel (gray).

The Q torus has high net positive charge (+4, excluding sidechains that interact with RNAP), consistent with accommodation of negatively-charged RNA. Protein-RNA interactions by the Q torus involve predominantly the RNA backbone, and, at the narrowest point of the Q torus, protein-RNA interactions by the Q torus involve exclusively the RNA backbone (Figs. 3D and S6C). At the narrowest point of the Q torus, the sidechains of Q21 Ser29 and Thr54 are positioned to form H-bonds with a phosphate of the RNA backbone (Fig. 3D) and are positioned such that rotation about the Cα-Cβ bond could allow each to maintain the H-bond with a phosphate over ∼3 Å distance as a phosphate threads into and through the narrowest point, potentially providing a “molecular bearing” facilitating threading of RNA through the narrowest point. The presence of high net positive charge, the presence of protein-RNA interactions involving predominantly the RNA backbone, and the presence of a potential “molecular bearing,” provide a plausible basis for sequence-independent, sequence-nonspecific RNA threading though the Q torus during transcription elongation.

Comparison of the structures of the Q21-loading complex and the Q21-loaded complex reveals the mechanism for transformation of the loading complex to the loaded complex (Fig. 4A). The structures indicate the transformation involves five reactions: (i) threading of the RNA 5’ end into and through the Q torus, resulting in steric clash between the RNA 5’ end and the QBE; (ii) disruption of protein-DNA interactions by Qu (the Q21 protomer that makes weaker protein-DNA interactions and stronger protein-RNAP interactions); (iii) disruption of protein-RNAP interactions by Qd (the Q21 protomer that makes stronger protein-DNA interactions and weaker protein-RNAP interactions); (iv) disruption of σ-DNA and σ-RNAP interactions, and (v) rotation of the DNA-helix axis of upstream double-stranded DNA by ∼90° (purple, dark green, light blue, brown, and gray arrows in Fig. 4A). The structures explain why Q21 loading requires a promoter-proximally-paused TEC: namely, only a promoter-proximally-paused TEC has an accessible binding sites for Q21 on DNA (i.e., QBE, which becomes accessible only after promoter clearance; Fig. 1B, lines 2-3), has an accessible binding site for Q21 on RNAP (i.e., RNAP dock, which becomes accessible only after promoter clearance; Fig. 1B, lines 2-3), and has an RNA product short enough to thread into and through the Q torus, enabling establishment of topological linkage between the RNA product and the Q torus.

**Fig. 4.**
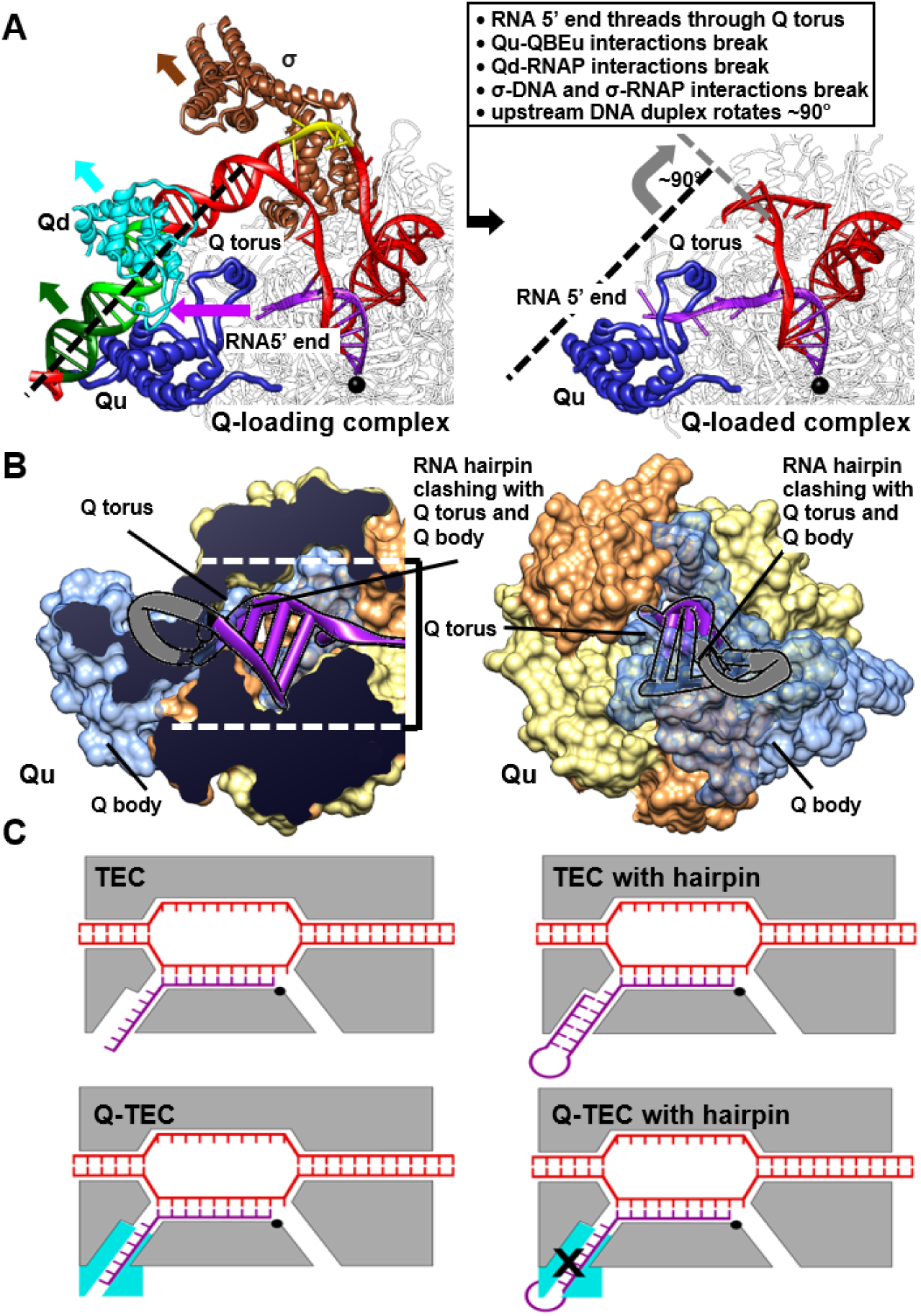
Mechanistic conclusions. **(A)** Transformation of Q21-loading complex to Q21-loaded complex. Black and gray dashed lines, DNA-helix axes of upstream dsDNA segments in loading and loaded complexes, respectively; purple, green, cyan, brown, and gray arrows, structural changes upon transformation of loading complex to loaded complex. Other colors as in Fig. 2B. **(B)** Antitermination and antipausing by Q21. Top, steric incompatibility of Q torus and pause and terminator RNA hairpins (hairpin stem from PDB 6ASX in purple, with segments positioned to interpenetrate Q-torus and Q body indicated as transparent ribbons with black outlines; hairpin loop from PDB 1MT4, which is positioned fully to interpenetrate Q-torus and Q body, indicated as gray ribbon with black outlines). View orientations and colors as in Figs. 2G and 3C. Bottom, schematic comparison of TECs in absence of Q (upper row) and TECs presence of Q (lower row). Colors as in Fig. 2B and 3B.

The structure of the Q21-loaded complex also reveals the mechanisms of antipausing and antitermination by Q21 (Fig. 4B). Hairpin-dependent pausing involves nucleation of an RNA hairpin at the mouth of the RNAP RNA-exit channel, followed by propagation of the RNA hairpin stem and penetration of the RNA hairpin stem into the RNAP RNA-exit channel, where the RNA hairpin stem makes RNA-RNAP interactions that promote pausing (Fig. 4B, top; 25–27). RNA hairpin-dependent transcription termination--the mode of termination that is most common in bacterial and bacteriophage transcription, and the mode of termination that occurs at the terminator preceding lambdoid bacteriophage late genes--involves nucleation of an RNA hairpin at the mouth of the RNA-exit channel, followed by propagation of the RNA hairpin stem and penetration of the RNA hairpin stem into the RNAP RNA-exit channel, followed by further propagation of the RNA hairpin stem that exerts mechanical forces that extract the RNA from RNAP or cause RNAP to translocate forward off the RNA (28). The structure of the Q21-loaded complex shows that Q21 is positioned to constitute a steric barrier to nucleation, propagation, and penetration of the RNA-exit channel by a pause or terminator hairpin (Fig. 4B). The Q body is positioned to overlap--essentially *in toto*--the loop and loop-proximal stem of a pause or terminator hairpin, and thus is positioned to block nucleation of a hairpin (Fig. 4B). The Q torus is positioned to overlap--*in toto* for the stem position in the narrowest point of the Q torus, and in part for the stem positions downstream of the narrowest point of the Q torus--one strand of the stem of a pause or terminator hairpin, and thus is positioned to block propagation and penetration of RNA-exit channel by a hairpin (Fig. 4B). Most important, the Q torus, with dimensions that accommodate only one strand of RNA constitutes an effectively absolute steric barrier to double-stranded RNA secondary structure at the mouth of, and inside, the RNAP RNA-exit channel (Fig.4B).

The structures of the Q-loading and Q-loaded complexes suggest that, in addition, Q21 may exert antipausing activity by inhibiting RNAP “swivelling”--an ∼3° rotation of an RNAP “swivel module,” comprising the RNAP dock, zinc binding domain (ZBD), lid, clamp, shelf, jaw, and SI3, that is thought to be associated with pausing (25-27,29-30; Fig. S7). Model building indicates that interactions of Q21 H2 with the RNAP ZBD and RNAP lid inside the RNAP RNA-exit channel could sterically preclude both the ∼3° swivelling associated with hairpin-dependent pausing (25-27) and the smaller, ∼1.5°, swivelling associated with elemental pausing (25; referred to as “pre-swivelling” in *30*; Fig. S7A-B). Specifically, model building indicates that swivelling would increase distance and break interactions between Q H2 and the RNAP ZBD and, simultaneously, would decrease distance and create steric clash between H2 and the RNAP lid (Fig. S7A-B). The steric incompatibility of Q21 to swivelling may contribute to Q21 antipausing, and, in particular, may provide the mechanism to trigger the release of the Q-loading complex from the SDPE.

Taken together, the results presented here reveal that Q21 forms a torus--a molecular “nozzle”-- that narrows and extends the RNAP RNA exit channel, that the RNA product is threaded through the Q nozzle, and that RNA threading through the Q nozzle prevents the formation of pausing and terminator hairpins and the prevents RNAP swivelled state associated with pausing (Figs. 4B and S7B).

Narrowing and extending the RNAP RNA-exit channel by attaching a molecular nozzle is a strikingly simple way to achieve antitermination--much simpler than the recently elucidated mechanism of the other textbook example of transcription antitermination factor, lambdoid bacteriophage N (31)--and almost surely will be a generalizable mechanism.

Functional analyses and sequence analyses have identified more than 15,000 lambdoid bacteriophage Q proteins, comprising three protein families: the Q21/Q933W family (Pfam PF06530; 5,251 entries in NCBI), the Qλ family (Pfam PF03589, 7,904 entries in NCBI), and the Q82 family (Pfam PF06530, 2,635 entries in NCBI). Q proteins from the three protein families perform equivalent regulatory functions and are encoded by genes that exhibit equivalent positions in bacteriophage genomes, but, surprisingly, Q proteins from the three protein families exhibit no significant sequence similarity to each other (11,14-15,32). Our crystal structure of Q21 (Fig. S1) shows no significant three-dimensional structural similarity to a previously reported crystal structure of a fragment of Qλ (33), showing that the lack of sequence similarity is accompanied by a lack of three-dimensional structural similarity. A key priority for further research is to determine whether the functionally equivalent, but non-homologous, Q proteins of the Qλ and Q82 protein families also function by narrowing and extending the RNAP RNA-exit channel by attaching a molecular nozzle, and, if so, how they use non-homologous sequences and three-dimensional structures to do so.

Another striking aspect of the molecular-nozzle mechanism of Q21 is that the mechanism results in a topological linkage of Q21 and the RNA product (Figs. 4 and S6), establishing an unbreakable linkage between Q21 and the TEC that enables Q21 to provide processive antipausing and antitermination activity over tens of thousands of nucleotide-addition steps. A key priority for future research is to determine whether Q21 can function with--or can be engineered to function with--TECs from other species, enabling this exceptional topological linkage of a regulatory factor and a TEC, and the resulting exceptionally processive antipausing and antitermination activity, to be broadly exploited for regulated gene expression, synthetic biology, and metabolic engineering.

## Acknowledgements

This work was supported by National Institutes of Health grant GM041376 to R.H.E.. We thank the Argonne National Laboratory and the National Synchrotron Light Source II at Brookhaven National Laboratory for beamline access; the Rutgers University Cryo-EM Core facility and the John M. Cowley Center for High Resolution Electron Microscopy at Arizona State University for microscope access; I. Artsimovitch, S. Borukhov, and B. Nickels for plasmids; B. Nickels and T. Santangelo for discussion; and Williams for assistance.

## Supplementary Materials

### Materials and Methods

#### Bacteriophage 21 Q (Q21)

*E. coli* strain BL21 Star (DE3) (Invitrogen) was transformed with plasmid pET21-H6-TEV-Q21, encoding N-hexahistidine-tagged, tobacco-etch-virus-protease-site-tagged Q21 under control of the bacteriophage T7 gene 10 promoter [constructed by replacing the NdeI-SalI segment of plasmid pET21a (EMD Millipore) with an NdeI-SalI segment of a synthetic DNA fragment (Genscript) carrying 5’-CATATGGGACATCACCATCACCATCACGAGAACCTGTACTTCCAATCC-3’, followed by codons 2-162 of gene Q of lambdoid bacteriophage 21 (14), followed by 5’-GAGTCGAC-3’].

Single colonies of the resulting transformants were used to inoculate 200 ml LB broth containing 100 µg/ml ampicillin, and cultures were incubated 16 h at 37°C with shaking. Aliquots (20 aliquots of 15 ml each) were used to inoculate batches (20 batches of 1 L each LB broth containing 100 µg/ml ampicillin, cultures were incubated at 37°C with shaking until OD_600_ = 0.6, cultures were induced by addition of IPTG to 0.5 mM, and cultures were incubated overnight at 16°C. Cells from batches were harvested by centrifugation (4,000 × g; 15 min at 4°C), were pooled and re-suspended in 200 ml buffer A [20 mM Tris-HCl, pH 8.0, 1 M NaCl, 0.5 M NaBr, 20 mM imidazole, and 2 mM tris(2-carboxyethyl)phosphine (TCEP; Sigma Aldrich)], and were lysed by sonication [Branson Sonifier 450 (VWR); 0.5 s bursts at 0.5 s intervals at output setting 10 over 30 min on ice]. The lysate was centrifuged (13,000 × g; 30 min at 4°C), the supernatant was loaded onto a 10 ml column of Ni-NTA-agarose (Invitrogen) equilibrated in buffer A, and the column was washed with 100 ml buffer A and eluted with 200 ml a linear gradient of buffer A to buffer A containing 500 mM imidazole. Fractions containing Q21 ((fractions eluting at ∼200 mM imidazole; 50 ml total) were collected, pooled, were supplemented with 1 ml (4 mg/ml) hexahistidine-tagged tobacco-etch-virus protease (prepared as in 34) in buffer A, incubated with simultaneous dialysis against 100 volumes buffer B (20 mM Tris-HCl, pH 8.0, 1 M NaCl, and 2 mM TCEP) for 16 h at 6°C?, to convert tagged Q21 to untagged Q21, were further purified by applying to a 10 ml column of Ni-NTA-agarose equilibrated in buffer B and collecting the flow-through, were further purified by size-exclusion chromatography on a Superdex S200 26/60 column (GE Healthcare) equilibrated and eluted in buffer C (10 mM Tris-HCl, pH 7.4, 100 mM NaCl, and 1 mM dithiothreitol), were concentrated to 8 mg/ml in buffer C using 10 kDa MWCO Amicon Ultra-15 centrifugal ultrafilters (Millipore), and were stored at −80°C. Yields were ∼2.5 mg/L, and purities were >95%.

Selenomethionine (SeMet)-substituted Q21 produced using SeMet Minimal Medium (Molecular Dimensions) following the procedures of the manufacturer and was purified as for native Q21.

#### E. coli GreB

N-terminally hexahistidine-tagged *E. coli* GreB was prepared using *E. coli* strain XL-1 Blue (Agilent) transformed with plasmid pTRC-GreB-NT-His (35), as described (35).

#### *E. coli* [Cys541;Pro607]-σ^70^

N-terminally hexahistidine-tagged [Cys541;Pro607]-σ^70^ (*E. coli σ*^70^ derivative with reduced affinity for RNAP FTH, resulting in increased proficiency in Q loading; *8,10,36*) was prepared using *E. coli* strain BL21 Star (DE3) (Invitrogen) transformed with plasmid pLHN12-His (36), as described (36).

#### *E. coli* RNA polymerase core enzyme

*E. coli* RNAP core enzyme containing C-terminally decahistidine-tagged β’ subunit was prepared using *E. coli* strain BL21 (DE3) (Invitrogen) transformed with plasmid pIA900 (encodes *E. coli* RNAP β’, β, α, and ω subunits under control of the bacteriophage T7 gene 10 promoter; *37*), as described (37).

#### *E. coli* RNA polymerase σ^70^ holoenzyme

*E. coli* RNAP σ^70^ holoenzyme containing C-terminally decahistidine-tagged β’ subunit was prepared using *E. coli* strain BL21 (DE3) (Invitrogen) co-transformed with plasmids pIA900 [encodes *E. coli* RNAP β’, β, α, and ω subunits under control of the bacteriophage T7 gene 10 promoter; *37*] and pRSF-Duet-Ec-rpoD [encodes *E.* coli σ^70^; constructed by replacing the NcoI-HindIII segment of plasmid pRSF-Duet (EMD Millipore) with an NcoI-HindIII segment of a PCR-generated DNA fragment carrying 5’- CCATGGG-3’, followed by codons 1-613 of the *E. coli rpoD* gene (38), followed by 5’- TAACACGACAAGCTT-3’], as described (37).

#### Oligonucleotides

Oligodeoxyribonucleotides (IDT; fluorescein-labelled oligodeoxyribonucleotides synthesized using 5’ 6-FAM; IDT), and oligoribonucleotides (IDT) were dissolved in nuclease-free water (Ambion.) to 1 mM and stored at −80°C.

#### DNA fragments and nucleic-acid scaffolds

DNA fragments for preparation and analysis of Q21-QBE complexes (sequences in Figs. 1C and S2D) were prepared as follows: Nontemplate-strand oligodeoxyribonucleotide (0.5 mM), and template-strand oligodeoxyribonucleotide (0.5 mM) in 0.5 500 µl 10 mM Tris-HCl, pH 8.0, and 50 mM NaCl were heated 10 min at 95°C, slow-cooled to 25°C (∼2 h), and stored at −80°C.

The nucleic-acid scaffold for preparation and analysis of the Q21-loading complex (sequence in Fig. 1C and SA) was prepared as follows: Nontemplate-strand oligodeoxyribonucleotide (0.3 mM), template-strand oligodeoxyribonucleotide (0.3 mM), and oligoribonucleotide (0.33 mM) in 500 µl 10 mM Tris-HCl, pH 8.0, and 50 mM NaCl were heated to 95°C for 2.5 min, slow-cooled to 25°C (∼3 h) and then to 4°C (∼0.5 h), and stored at −80°C.

The DNA fragment for preparation and analysis of the Q21-loaded complex (sequence in Fig. S5A) was generated by PCR amplification (39) using a full-length nontemplate-strand oligodeoxyribonucleotide as template and a 18 nt 5’ nontemplate-strand oligodeoxyribonucleotide and 18 nt 5’ template-strand oligodeoxyribonucleotide as primers; was purified by electrophoresis on 1.5 % agarose slab gels (Sigma Aldrich) in TAE buffer (39), followed by excision of gel slices, and extraction from gel slices using the QIAquick Gel Extraction Kit (Qiagen) according to the instructions of the manufacturer.

#### Protein-DNA interaction assays: fluorescence polarization

Equilibrium fluorescence polarization assays of protein-DNA interaction (40) were performed in a 96-well microplate format. Reaction mixtures (200 µl) containing 0 or 2 μM Q21 in buffer C were serially diluted (2-fold serial dilutions) with 10 nM fluorescein-labelled DNA fragment in 20 mM Tris-HCl pH 8, 50 mM NaCl, 1 mM DTT and 0.02% Triton-X, equilibrating 10 min at 25°C following each serial dilution, in black 96-well polystyrene microplates (catalog number 655209; Greiner Bio-One). Fluorescence emission intensities were measured using a microplate reader equipped with polarization filters (GENios Pro; TECAN; excitation wavelength = 485 nm; emission wavelength = 535 nm). Emission intensities were corrected for background by subtracting emission intensities in titrations with 0 μM Q21, fluorescence polarization was computed using Magellan Data Analysis Software (TECAN), and data were fit to a mathematical model of one-site specific binding with ligand depletion (41) using Prism v5.01 (GraphPad Software).

#### Crystal structure determination: assembly of Q-QBE complex

The Q21-QBE complex was prepared by incubating 500 μM Q21 and 250 μM QBE DNA fragment in 2 ml buffer C for 10 min at 2°C5, was purified by size-exclusion chromatography on a Superdex S200 26/60 column (GE Healthcare, Inc) equilibrated and eluted in buffer C, was concentrated to 6 mg/ml in buffer C using 10 kDa MWCO Amicon Ultra-15 centrifugal ultrafilters (Millipore), and was stored at −80°C.

#### Crystal structure determination: crystallization and data collection

Robotic crystallization trials were performed for Q21 and Q21-QBE using a Gryphon liquid handling system (Art Robbins Instruments,), commercial screening solutions (Emerald Biosystems, Hampton Research, and Qiagen), and the sitting-drop vapor diffusion technique [drop: 0.5 µl μM Q21 or Q21-QBE in buffer C plus 0.5 µl screening solution; reservoir: 60 µl screening solution; 6°C or 16°C]. 900 conditions each were screened for Q21 and Q21-QBE, and conditions yielding crystals were optimized using the hanging-drop vapor-diffusion technique.

Crystals of native Q21 for X-ray diffraction data collection were prepared using the hanging-drop vapor diffusion method, by mixing 1 µl 400 µM Q21 in buffer C with 1 μl of a reservoir solution consisting of 20% (w/v) polyethylene glycol 3350 (Hampton Research) and 200 mM ammonium chloride at 6°C. Crystals of SeMet-derivatized Q21 for X-ray diffraction data collection were prepared using the hanging-drop vapor diffusion method by mixing 1 µl of 400 μM SeMet-derivatized Q21 in buffer C with 1 μl of a reservoir solution consisting of 20% (w/v) polyethylene glycol 8000 (Hampton Research) and 100 mM Hepes-NaOH, pH 7.5, at 6°C. Crystals of native Q21 and SeMet-derivatized Q21 were cryo-protected by transfer to 5 μl reservoir solution containing 15% ethylene glycol and were cooled by immersion in liquid nitrogen. X-ray diffraction data were collected on single crystals of native Q21 and SeMet-derivatized Q21 at 100°K on at the Argonne National Laboratory Advanced Photon Source beamline 19-ID (wavelength 0.987 Å).

Crystals of Q21-QBE for X-ray diffraction data collection were prepared with the sitting-drop vapor diffusion method by mixing 1 µl of 320 µM Q21-QBE in buffer C with 1 µl reservoir solution consisting of 0.2 M NaCl, 0.1 M bis-Tris-HCl, pH 6.5, and 25% (w/v) polyethylene glycol 3350. Crystals of Q21-QBE were cryo-protected by transfer to 5 μl reservoir solution containing 15% ethylene glycol and were cooled by immersion in liquid nitrogen. X-ray diffraction data were collected on a single crystal of Q21-QBE 100°K at the Brookhaven National Laboratory National Synchrotron Light Source II beamline 17-ID-1 (wavelength 0.97 Å).

#### Crystal structure determination: structure solution and refinement

Data were processed, and integrated intensities were merged and scaled, with automatic correction and a resolution cut-off of I/σ = 2.0, using HKL2000 (42).

The crystal structure of SeMet-derivatized Q21 was solved by experimental phasing with single-wavelength anomalous dispersion (43) in Phenix (44), using anomalous diffraction data collected on a crystal of SeMet-derivatized Q21. Thirteen heavy-atom sites were identified, and an initial model was built in Phenix. The crystal structure of native Q21 was solved by molecular replacement (45) in Phenix, using the crystal structure of SeMet-derivatized Q21 as the search model. The crystal structure of Q21-QBE was solved by molecular replacement in Phenix, using the crystal structure of native Q21 as the search model. Iterative cycles of model building in Coot (46) and refinement in Phenix then were performed, adding riding hydrogen atoms to all models, performing translation-libration-screw-rotation parameterization by TLSMD analysis (47), and performing validation with MolProbity (48).

Final models of SeMet-derivatized Q21 (residues 6-107, 110-114, and 117-159) refined to 2.0 Å, native Q21 (6-156) refined to 1.94 Å, and Q21-QBE (Q21 residues 5-156, nontemplate-strand nucleotides 1-21, and template-strand nucleotides 1-21) refined to 2.84 Å, have been deposited in the Protein Data Bank (PDB) with accession codes 6P1C, 6P1B, and 6P1A, respectively (Table S1).

#### Cryo-EM structure determination (Q21-loading complex): sample preparation

The Q21-loading complex was prepared by incubating 8 μM *E. coli* RNAP core enzyme with 10 μM nucleic-acid scaffold (sequence in Fig. 2A) in 1 ml buffer D (10 mM Tris-HCl, pH 7.8, 100 mM NaCl, 1 mM MgCl_2_, and 1 mM dithiothreitol) 5 min at 25°C, followed by adding 400 μl 28 μM *E. coli* [C541; P607] σ^70^ in buffer D and 200 μl 220 μM Q21 in buffer D and incubating 5 min at 25°C. The sample was purified by size-exclusion chromatography on a Superdex S200 26/60 column (GE Healthcare) equilibrated and eluted in buffer D, was concentrated to 9 mg/ml in buffer D using 10 kDa MWCO Amicon Ultra-15 centrifugal ultrafilters (Millipore), and was stored at −80°C.

EM grids were prepared using a Vitrobot Mark IV autoplunger (FEI), with the environmental chamber at 22°C and 100% relative humidity. A sample comprising 45 μl 20 μM Q21-loading complex in buffer D was mixed with 5 μl 80 mM CHAPSO (Hampton Research) in water 5 min on ice. Aliquots (2.5 μl) were applied to glow-discharged (5 min) UltrAuFoil (1.2/1.3) 300-mesh grids (Quantifoil), grids were blotted with Standard Vitrobot Filter Paper (Agar Scientific) 3 s at 22°C, and grids were flash-frozen by plunging in a liquid ethane cooled with liquid nitrogen, and grids were stored in liquid nitrogen.

#### Cryo-EM structure determination (Q21-loading complex): data collection and data reduction

Cryo-EM data were collected at the John M. Cowley Center for High Resolution Electron Microscopy at Arizona State University, using a 300 kV Titan Kiosk (FEI/ThermoFisher) electron microscope equipped with a K2 Summit direct electron detector (Gatan). Data were collected automatically in super-resolution counting mode using SerialEM (49), a nominal magnification of 22,500x (actual magnification 47,608x), a calibrated pixel size of 0.553 Å per super-resolution pixel, and a dose rate of 8 electrons/pixel/s. Movies were recorded at 200 ms/frame for 6 s (30 frames total), resulting in a total radiation dose of 41.6 electrons/Å^2^. Defocus range was varied between −0.80 µm and −2.4 µm. A total of 3,010 micrographs were recorded from one grid over four days. Micrographs were saved as 4-bit LZW compressed Tiff images and were decompressed to 8-bit MRC upon pre-processing for gain normalization and defect correction.

Data were processed as summarized in Figs. S3-S4. Data processing was performed on a Tensor TS4 Linux GPU workstation (Exxact) containing four GTX 1080 Ti graphic cards (Nvidia). Dose weighting, motion correction (5×5 tiles; b-factor = 150), and Fourier-binning (2×2; pixel size = 1.066 Å) were performed using MotionCor2 (50). Contrast-transfer-function (CTF) estimation was performed using Gctf (51). Subsequent image processing was performed using Relion 3.0 (52). Automatic particle picking with Laplacian-of-Gaussian filtering yielded an initial set of 1,045,380 particles. Particles were extracted into 256×256 pixel boxes, 4x down-scaled, and subjected to rounds of reference-free 2D classification and removal of poorly populated classes, yielding a selected set of 529,871 particles. The selected set was 3D-classified with C1 symmetry, using a 60 Å low-pass-filtered map calculated from a cryo-EM structure of an *E. coli* TEC (PDB 6ALF; 53) as the 3D template, and the best-resolved class, comprising 254,610 particles, was 3D auto-refined and then subjected to 3D sub-classification without alignment, using the auto-refined map as the 3D template. Following the sub-classification, sub-classes exhibiting resolved Q21-QBE density were combined, and the resulting 150,310 particles were re-centered, re-extracted without down-scaling and subjected to 3D auto-refinement without a reference mask, using the previous auto-refined map as the 3D template. The resulting refined particles were CTF refined, Bayesian polished (54), and further CTF refined. The resulting CTF-refined particles were refined using a soft mask and solvent flattening and were post-processed, yielding a reconstruction at 3.5 Å overall resolution, as determined from gold-standard Fourier shell correlation (FSC; 55-56), using phase-randomization to account for convolution effects of a solvent mask on FSC between two independently refined half maps, “half-map 1” and “half-map 2” (57; Figs S3E, S4A; Table S2).

The initial atomic model for the Q21-loading complex was built by manual docking of the crystal structure of Q21-QBE (Fig. 1C-E, S2); RNAP β’, β, α^I,^, α^II^, and ω segments from a crystal structure of a *E. coli* transcription initiation complex (PDB 5IPL; 58); σ^70^ segments from a crystal structure of *E. coli* RNAP holoenzyme (PDB 4MEY; 59l); DNA segments from a crystal structure of an *E. coli* transcription initiation complex (PDB 4YLN; 60), and RNA segments from a cryo-EM structure of an *E. coli* TEC (PDB 6ALF; 53), using UCSF Chimera (61). For the Qu N-terminus (residues 1-6), the Qd N- and C-termini (residues 1-5 and 150-162), the RNAP β’ N and C-termini and SI3 (residues 1-14, 1376-1407, and 931-956), the RNAP β N-terminus (residues 1-2), the RNAP α^I^ N-terminus and C-terminal domain (residues 1-5 and 235-329), the RNAP α^II^ N-terminus and C-terminal domain (residues 1-5 and 233-329), the RNAP ω N- and C-termini (residues 1 and 76-91), σ^70^ region 1.1 (residues 1-89), parts of σR2 (residues 166-214, 238-241), and the C-terminal part of σR3 through the σR3-σR4 linker and σR4 (residues 467-613), density was absent, suggesting high segmental flexibility; these segments were not fitted.

Refinement of the initial model was performed using phenix.real_space_refine under Phenix (44,62-63). Each chain was rigid-body refined against the map, and then each chain was subjected to iterative cycles of model building and refinement in Coot (46), followed by real-space refinement with geometry, rotamer, Ramachandran-plot, Cβ, non-crystallographic-symmetry, secondary-structure, and reference-structure (initial model as reference) restraints, followed by global minimization and local rotamer fitting. Secondary-structure annotation was inspected and edited using UCSF Chimera, and geometry was monitored using MolProbity (44). Following model building for each chain, chains were combined and were further refined with geometry, rotamer, Ramachandran-plot, Cβ, non-crystallographic-symmetry, and secondary-structure restraints. The final refined model was validated against over-fitting by randomizing atomic coordinates by 0.3 Å; performing real-space refinement with geometry, rotamer, Ramachandran-plot, Cβ, non-crystallographic-symmetry, secondary-structure, and reference-structure (final refined model as reference) restraints against half-map 1; and performing FSC calculations comparing the resulting model to half-map 1, half-map 2, and the full map (Fig. S3E). Following validation of the final refined model, atom-displacement factors (*B*-factors) were real-space refined against the full map. The final atomic model has been deposited in the Electron Microscopy Data Bank (EMDB) and the PDB with accession 20233 and 6P18, respectively (Fig. S4B-C; Table S2).

#### Cryo-EM structure determination (Q21-loaded complex): sample preparation

The Q21-loaded complex was prepared in reaction mixtures containing (4 ml): 0.6 μM DNA fragment (sequence in Fig S5A), 0.6 μM *E. coli* RNAP σ^70^ holoenzyme, 4 μM *E. coli* GreB, 1.5 μM Q21, and 1 mM each of ATP, GTP, and UTP, in 10 mM Tris-HCl, pH 7.8, 50 mM KCl, 5 mM MgCl_2_, 1 mM dithiothreitol, and 5% glycerol. Reaction components except GreB, Q21, and the NTP subset were incubated 15 min at 37°C, GreB and Q21 were added and samples were incubated 1 min at 25°C, and the NTP subset was added and samples were incubated 15 min at 37°C. Products were purified by size-exclusion chromatography on a Superdex S200 26/60 column (GE Healthcare) equilibrated and eluted in buffer D; were concentrated to 8 mg/ml in buffer D using 10 kDa MWCO Amicon Ultra-15 centrifugal ultrafilters (Millipore); and were stored at −80°C. Electrophoresis of products on 7 M urea 10% polyacrylamide gels in TBE buffer (39), followed by staining with SYBR Green II RNA Gel Stain (Invitrogen) and UV imaging confirmed synthesis of a full-length, 70 nt, RNA product (Fig. S5B). Samples were applied to EM grids as described above for the Q21-loading complex.

#### Cryo-EM structure determination (Q21-loaded complex): data collection and data reduction

Cryo-EM data for the Q21-loaded complex were collected at the Rutgers University Cryo-EM Core Facility, using a 200 kV Talos Arctica (FEI/ThermoFisher) electron microscope equipped with a K2 Summit direct electron detector (Gatan) and a GIF Quantum imaging filter (Gatan) with slit width of 20 eV. Data were collected automatically in counting mode using EPU (FEI/ThermoFisher), a nominal magnification of 130,000x, a calibrated pixel size of 1.024 Å per pixel, and a dose rate of 6.4 electrons/pixel/s. Movies were recorded at 500 ms/frame for 5 s (25 frames total), resulting in a total radiation dose of 32.1 electrons/Å^2^. Defocus range was varied between −1.5 µm and −2.8 µm. A total of 1,528 micrographs were recorded from one grid over two days. Micrographs were gain-normalized and defect-corrected.

Data were processed as summarized in Figs. S5-S6. Dose weighting, motion correction, and CTF estimation were performed as described above for the Q21-loading complex. Automatic particle picking with Laplacian-of-Gaussian filtering yielded an initial set of 486,699 particles. Particles were extracted into 256×256 pixel boxes, 4x down-scaled, and subjected to rounds of reference-free 2D classification and removal of poorly populated classes, yielding a selected set of 218,536 particles. The selected set was 3D-classified with C1 symmetry using a 60 Å low-pass-filtered map calculated from a cryo-EM structure of an *E. coli* TEC (PDB 6ALF; 53) as the 3D template, and the best-resolved class, comprising 85,777 particles, was 3D auto-refined and then subjected to 3D classification without alignment, using the auto-refined map as the 3D template. Following the 3D-classification, the sub-class exhibiting resolved Q21 density was selected, and the 19,970 particles in the sub-class were re-centered, re-extracted without down-scaling and subjected to 3D auto-refinement without a reference mask, using the previous auto-refined map as the 3D-template. A focussed 3D classification without alignment then was performed using a custom mask generated by cropping density volume near the RNAP RNA-exit channel in Chimera and converting it to a soft-edged mask in Relion. From the focussed classification, sub-classes exhibiting consistent resolved density for Q21 were combined, the resulting 7,778 particles were refined without a mask, were CTF refined, Bayesian polished (54), and further CTF refined. The resulting CTF-refined particles were further refined using a soft mask and solvent flattening and were post-processed, yielding a reconstruction at 3.8 Å overall resolution, as determined from gold-standard FSC (55-56), using phase-randomization to account for convolution effects of a solvent mask on FSC between two independently refined half maps, “half-map 1” and “half-map 2” (57; Figs S5G; Table S3).

The initial atomic model for the Q21-loaded complex was built by manual docking of the cryo-EM structure of the Q21-loading complex (Figs. 2; S4; omitting atoms of Qd, σ^70^, DNA, and RNA), and DNA and RNA segments from a cryo-EM structure of *E. coli* TEC (PDB 6ALF; 53) and a crystal structure of a *Thermus thermophilus* TEC (PDB 2O5J; 64), using UCSF Chimera (61). For the Qu N-terminus and B1-B2 loop (residues 1-6 and 105-121), the RNAP β’ N and C-termini and SI3 (residues 1-14, 1376-1407, and 931-956), the RNAP β N-terminus (residues 1-2), the RNAP α^I^ N-terminus and C-terminal domain (residues 1-5 and 235-329), the RNAP α^II^ N-terminus and C-terminal domain (residues 1-5 and 233-329), the RNAP ω N- and C-termini (residues 1 and 76-91), σ^70^ region 1.1 (residues 1-89), parts of σR2 (residues 166-214, 238-241), and the C-terminal part of σR3 through the σR3-σR4 linker and σR4 (residues 467-613), density was absent, suggesting high segmental flexibility; these segments were not fitted.

Refinement and validation were performed as described above for the Q21-loading complex. The final atomic model was deposited in the EMDB and the PDB with accession codes 20234 and 6P19, respectively (Fig. S6; Table S3).

#### Structure analysis and molecular graphics

Molecular graphics representations were created using CCP4mg v2.9.0 (65) or UCSF Chimera (61). EM density maps were visualized in UCSF Chimera. Buried surface areas were determined using the PDBePISA server (66; https://www.ebi.ac.uk/pdbe/pisa/).

Structural similarity searches were performed using the DALI server (67; http://ekhidna2.biocenter.helsinki.fi/dali) and the PDBeFold server http://www.ebi.ac.uk/msd-srv/ssm).

## Supplementary Figure Legends

**Fig. S1.**
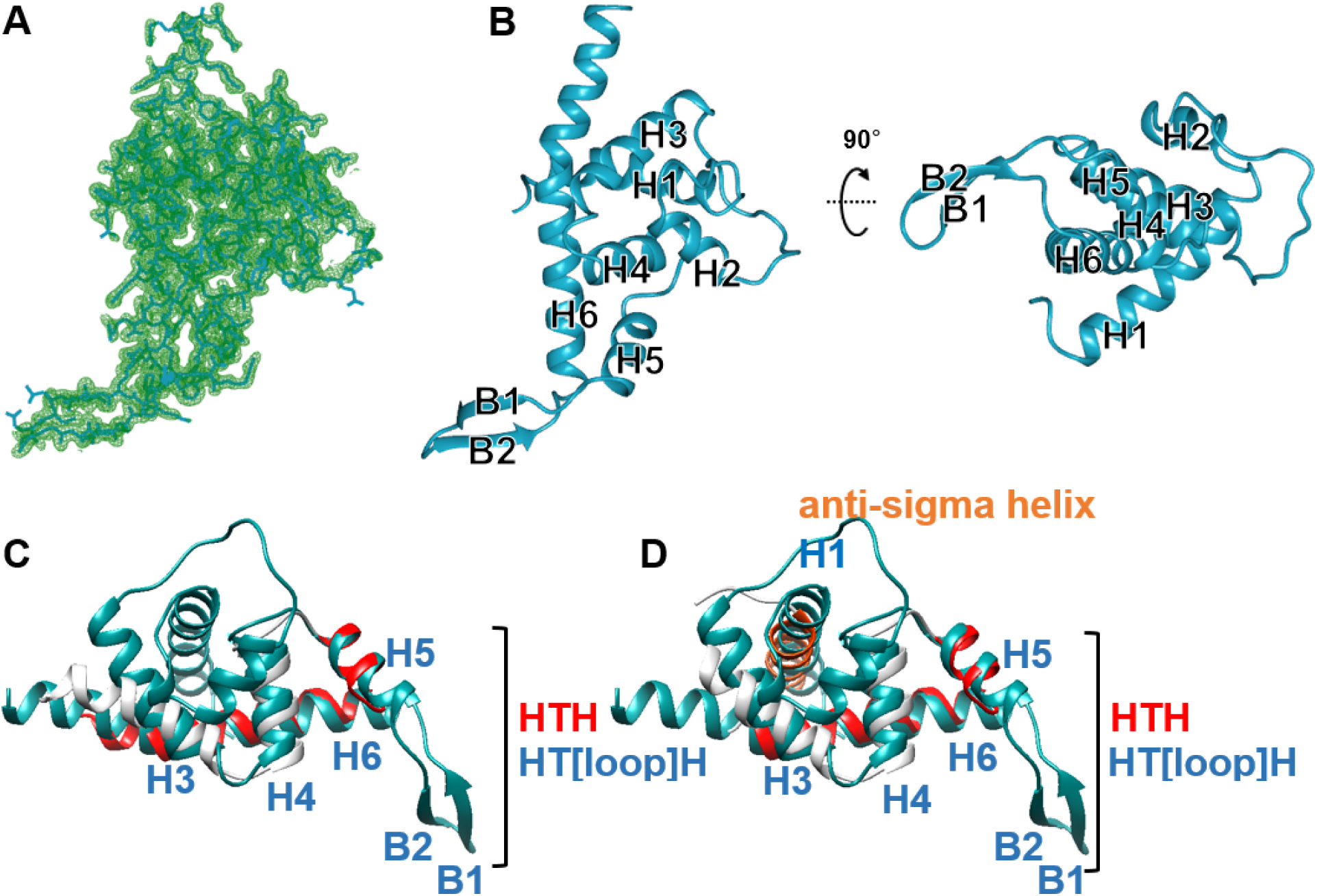
Crystal structure of Q21. **(A)** Crystal structure of Q21. Experimental electron density map (green mesh, 2mFo-DFc map contoured at 1.0σ) and fitted atomic model (blue). **(B)** Crystal structure of Q21 (ribbon representation; two orthogonal views). **(C)** Superimposition of Q21 (residues 7-156; blue) on ECF σ factor σR4 (PDB 2H27; 68; residues 122-190; gray, with HTH motif in red). DALI Z score = 6.4; PDBeFold Z score = 4.8. **(D)** Superimposition of Q21 (residues 7-156; blue) on ECF σ factor σR4 in complex with anti-sigma helix (PDB 1OR7; 69; σR4 residues 124-187 and anti-sigma residues 27-44; gray, with HTH motif in red and anti-sigma helix in orange). DALI Z score = 6.4; PDBeFold Z score = 5.5.

**Fig. S2.**
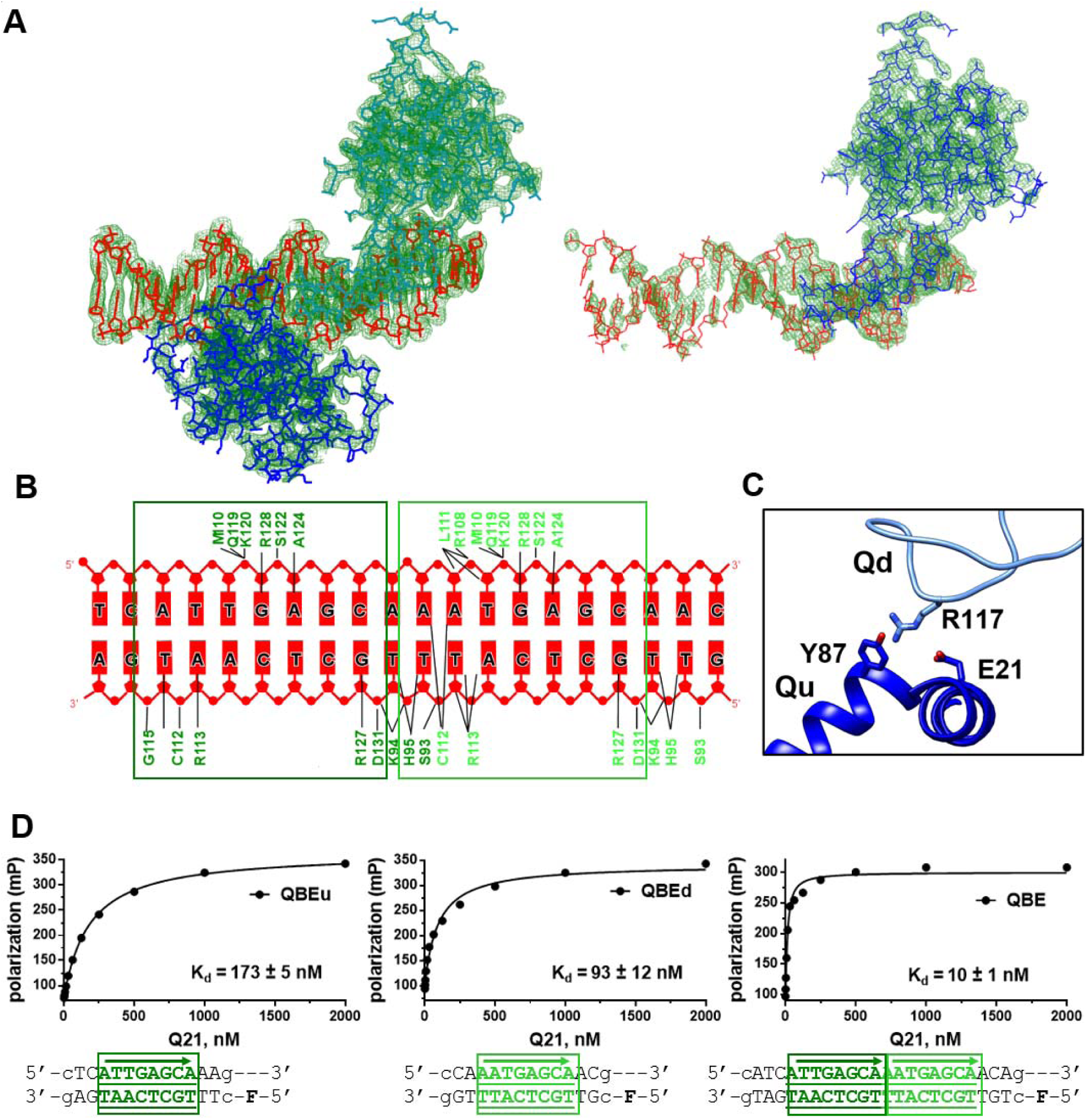
Crystal structure of Q21-QBE complex. **(A)** Crystal structure of Q21-QBE complex. Experimental electron density map (green mesh, 2mFo-DFc maps contoured at 1.0σ) and fitted atomic models (blue, light blue, and red) for the two molecular assemblies in the asymmetric unit: molecular assembly comprising two Q21 protomers bound to QBE (left) and molecular assembly comprising one Q21 protomer bound to QBE (right). **(B)** Summary of Q21-QBE interactions. Colors as in Fig. 1B. **(C)** Protein-protein interactions between Qd (B1-B2 loop of HT[loop]H motif; light blue) and Qu (H1 and H4; blue; sidechain oxygen atoms in red). **(D)** Fluorescence-polarization assays of Q21-DNA interactions with DNA fragments containing only QBEu (left), only QBEd (middle), and intact QBE comprising both QBEu and QBEd (right). Bold “F,” fluorescein.

**Fig. S3.**
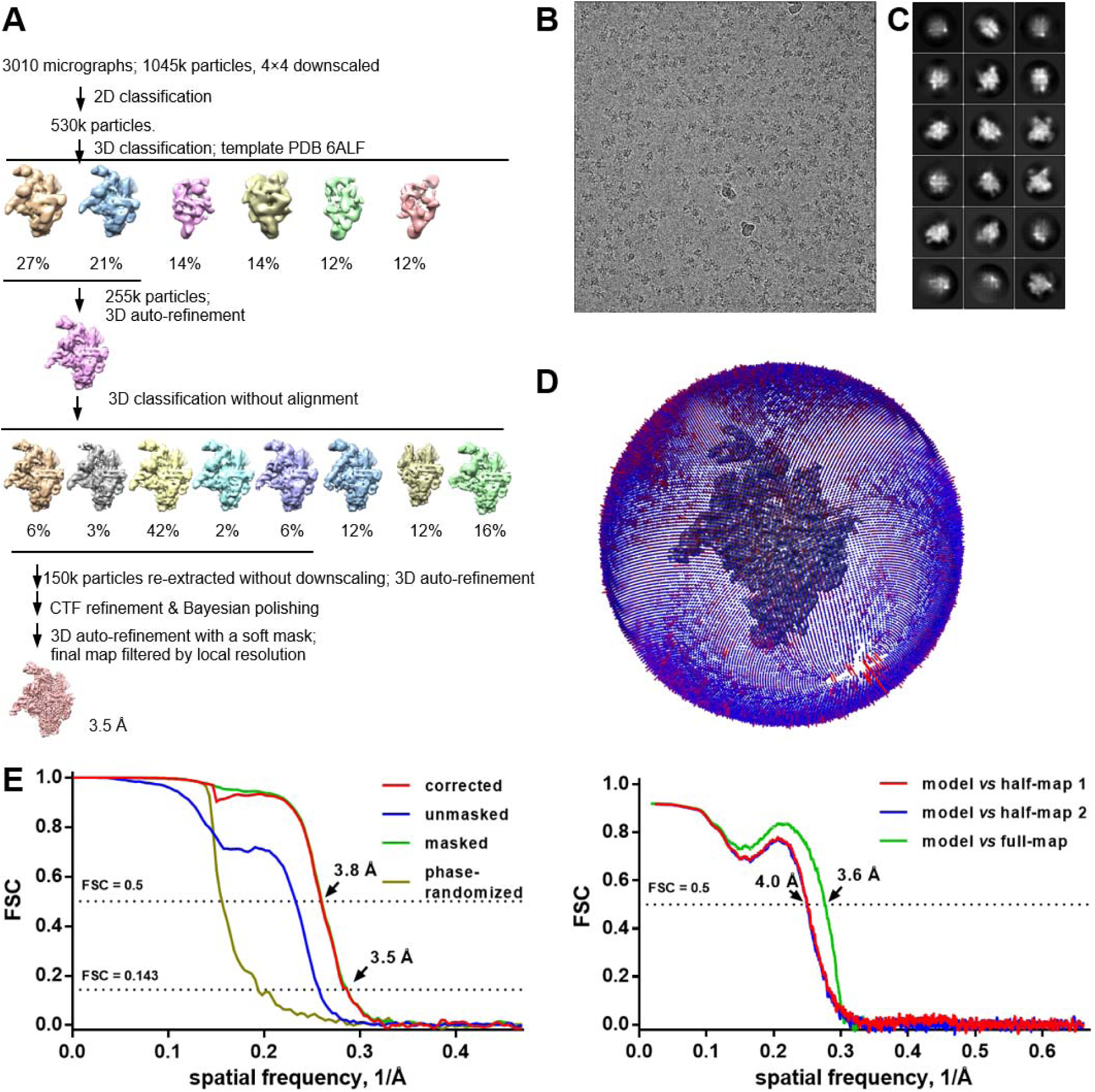
Cryo-EM structure of Q21-loading complex: structure determination. **(A)** Data processing scheme. **(B)** Representative electron micrograph. **(C)** 2D-class averages. **(D)** Euler-angle distribution. **(E)** Fourier-shell-correlation (FSC) plots.

**Fig. S4.**
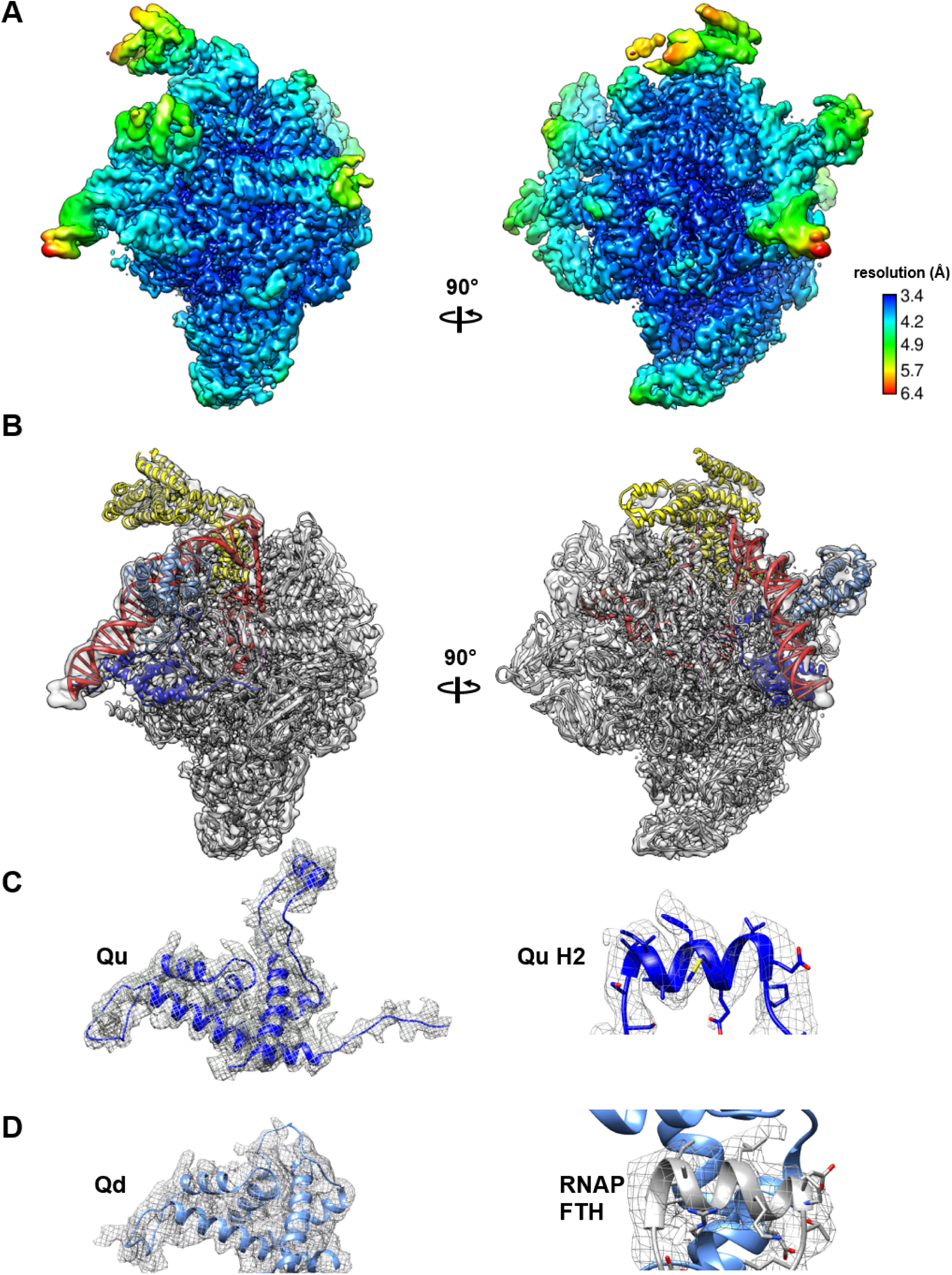
Cryo-EM structure of Q21-loading complex: maps and fit. **(A)** EM density map colored by local resolution. View orientation as in Fig. 2B. **(B)** EM-density map and fit. View orientation and colors as in Fig. 2B. **(C)** Density maps for Qu. Left, Qu. Right, close-up view of H2 of Q torus. **(D)** Density maps for Qd. Left, Qd. Right, close-up view of interaction between Qd and RNAP FTH.

**Fig. S5.**
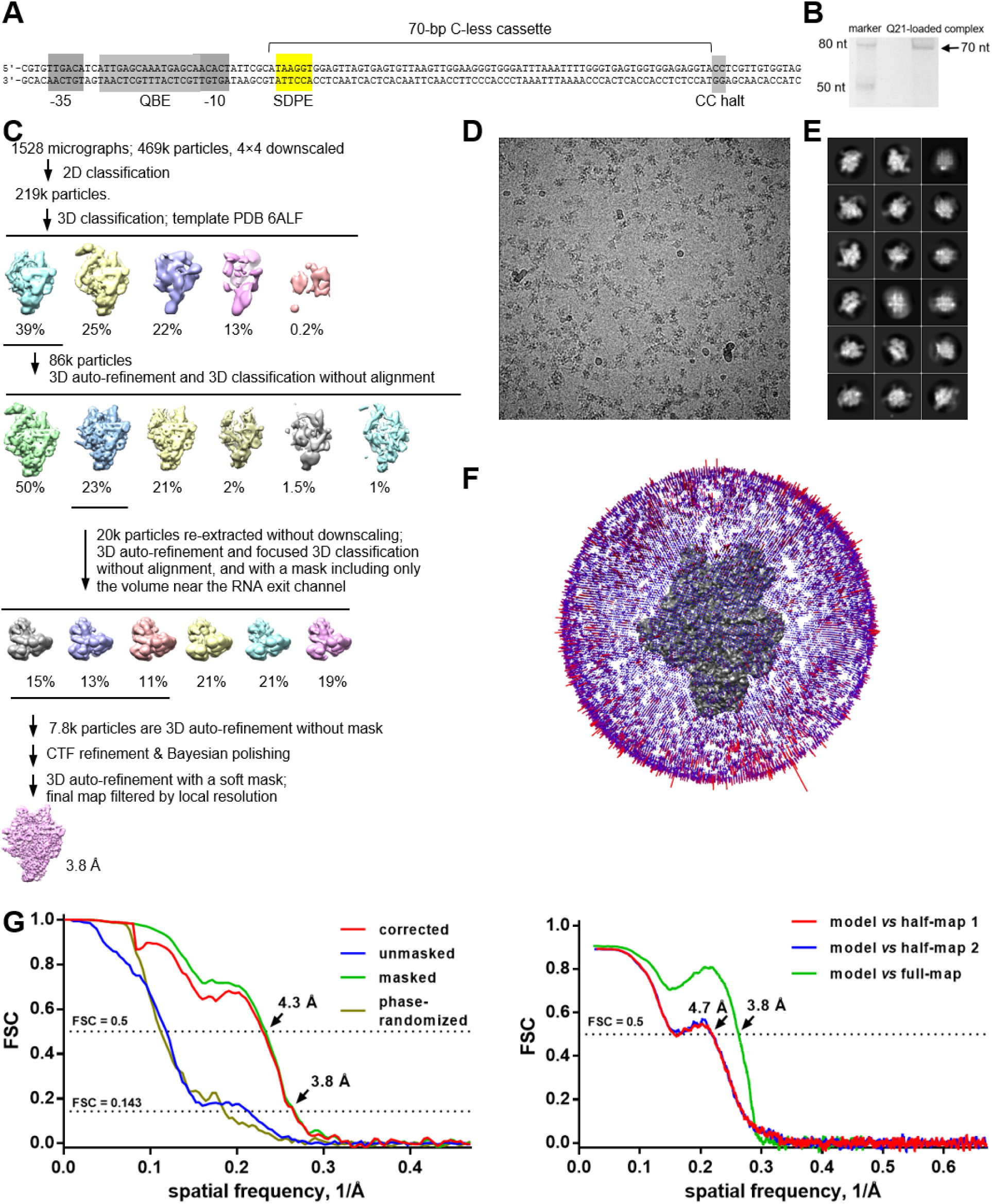
Cryo-EM structure of Q21-loaded complex: structure determination. **(A)** DNA template for preparation of Q21-loaded complex containing 70 nt RNA product.. **(B)** Preparation of Q21-loaded complex (urea-PAGE confirming generation of 70 nt RNA product). **(C)** Data processing scheme. **(D)** Representative electron micrograph. **(E)** 2D-class averages. **(F)** Euler-angle distribution. **(G)** Fourier-shell-correlation (FSC) plots.

**Fig. S6.**
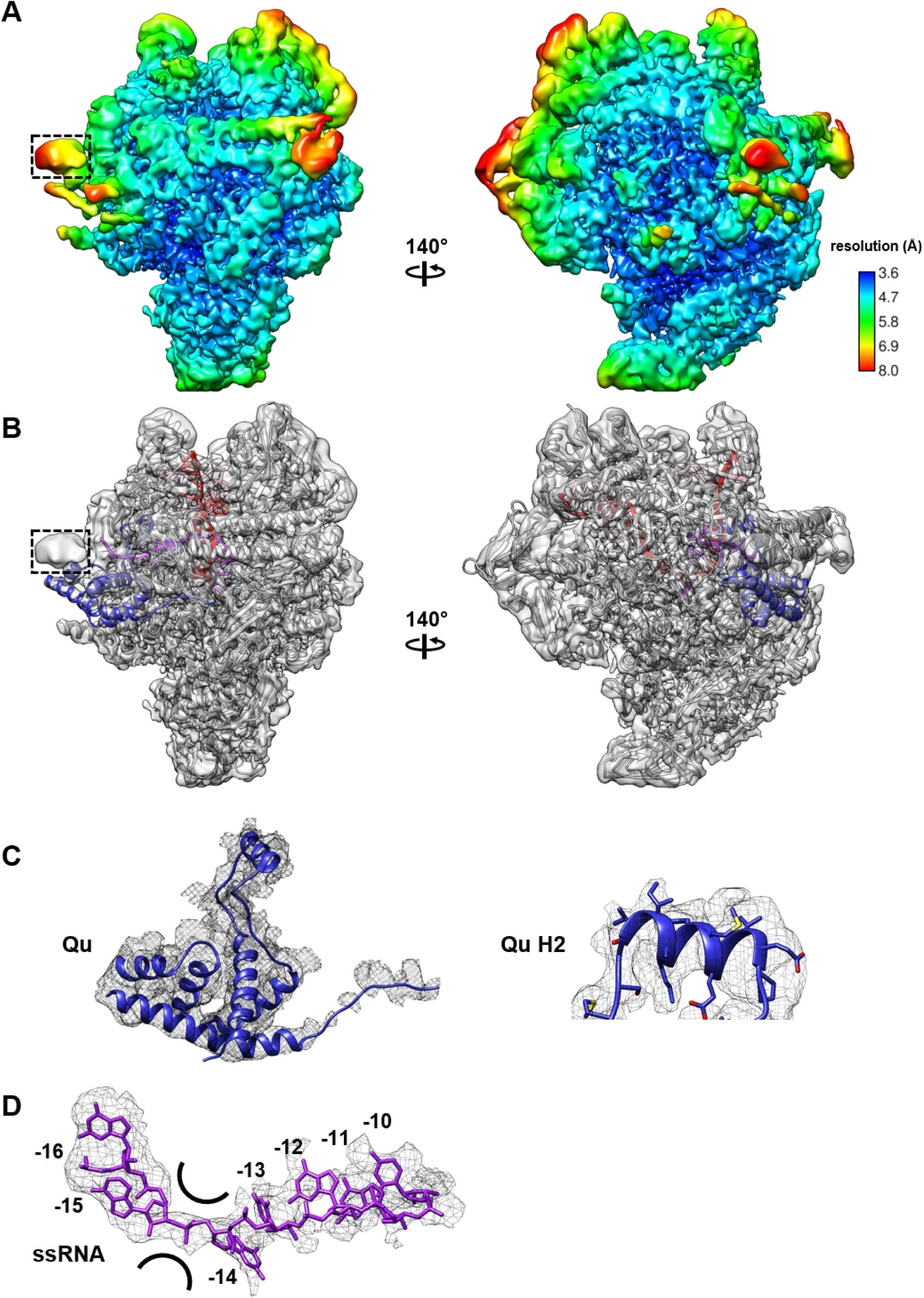
Cryo-EM structure of Q21-loaded complex: maps and fit. **(A)** EM density map colored by local resolution. Black dashed rectangle, unassigned density feature attributable to RNA nucleotides 5’ to RNA nucleotide −16 (RNA nucleotides numbered with RNA 3’ nucleotide as −1). View orientation as in Figs. 2B and 3B. **(B)** EM-density map and fit. View orientation and colors as in Figs. 2B and 3B. **(C)** Density maps for Qu. Left, Qu. Right, close-up view of H2 of Q torus. **(D)** Density map for RNA nucleotides −16 through −10 (numbered with RNA 3’ nucleotide as −1; narrowest position in Q torus indicated by black curved lines).

**Fig. S7.**
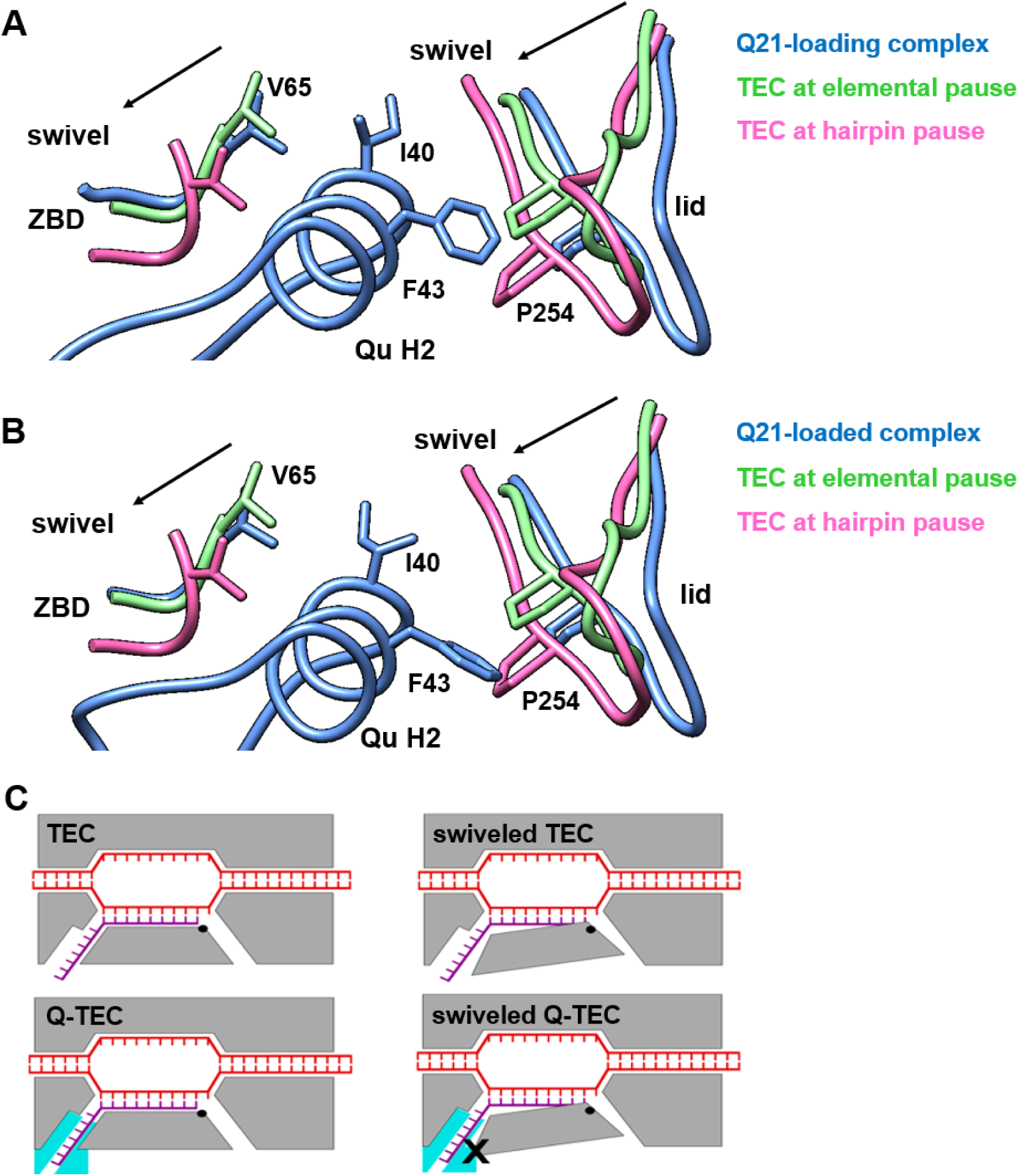
Suppression of RNAP swivelling by Q21. **(A)** Interactions of Q torus H2 with RNAP β’ zinc binding domain (ZBD) and RNAP β’ lid Q21-loading complex that preclude swivelling. Three structures are superimposed: Q-loading complex (blue), TEC with swivelling of ∼3°C associated with hairpin-dependent pausing (PDB 6ASX; 25; see also 26; pink), and TEC with swivelling of ∼1.5° associated with elemental pausing (PDB 6BJS; 25; green). Movement of RNAP ZBD and lid upon RNAP swivelling are indicated with arrows. RNAP swivelling is expected to increase distance and break interactions between Q H2 and the RNAP ZBD and, simultaneously, to decrease distance and create steric clash between H2 and the RNAP lid. **(B)** As A, but for Q21-loaded complex. **(C)** Schematic comparison of TECs in absence of Q (upper row) and TECs presence of Q (lower row). Colors as in Fig. 2B and 3B.

**Table S1.**
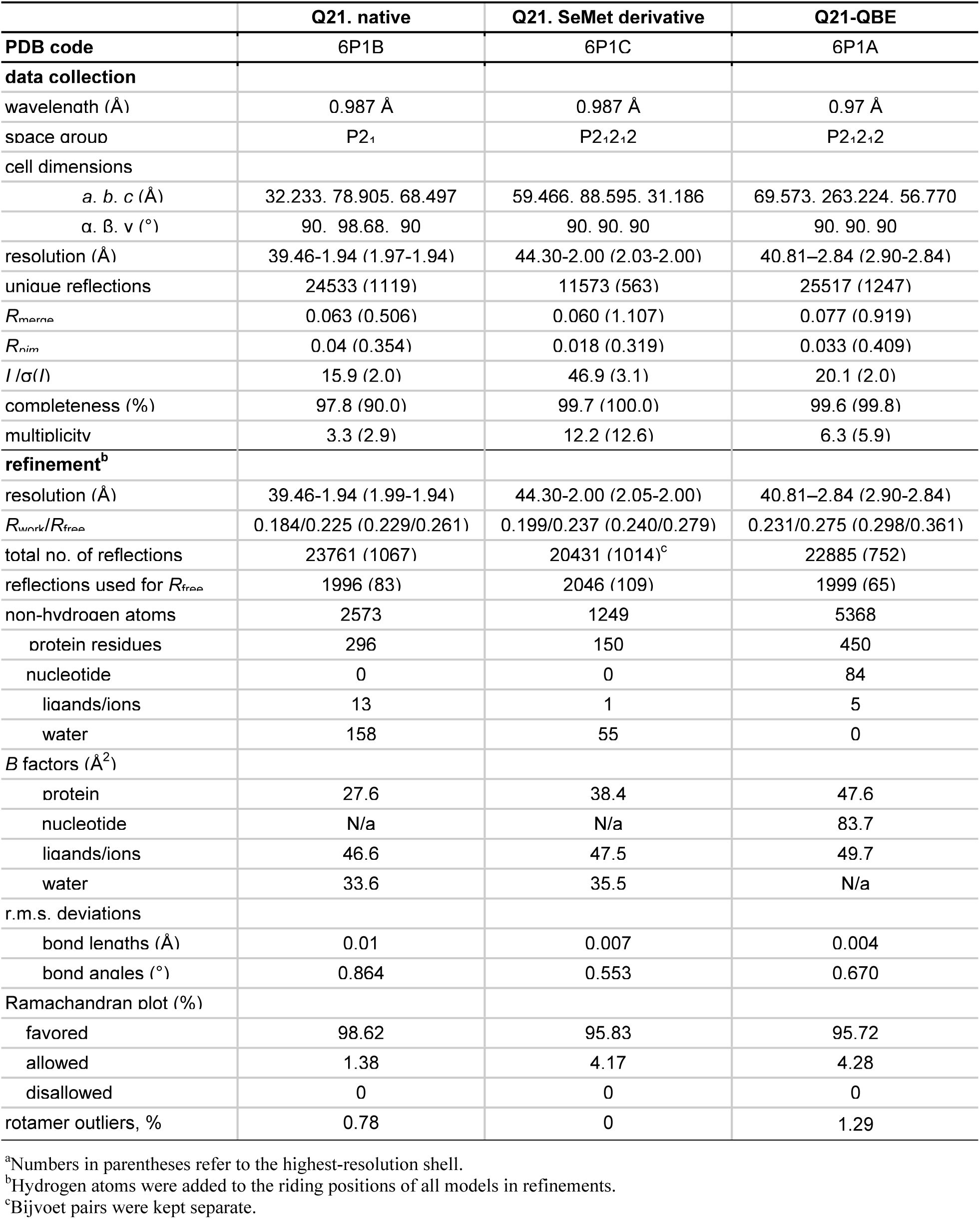
Data-collection and refinement statistics for crystal structures of Q21, SeMet-derivatized Q21, and Q21-QBE.^a^.

**Table S2.**
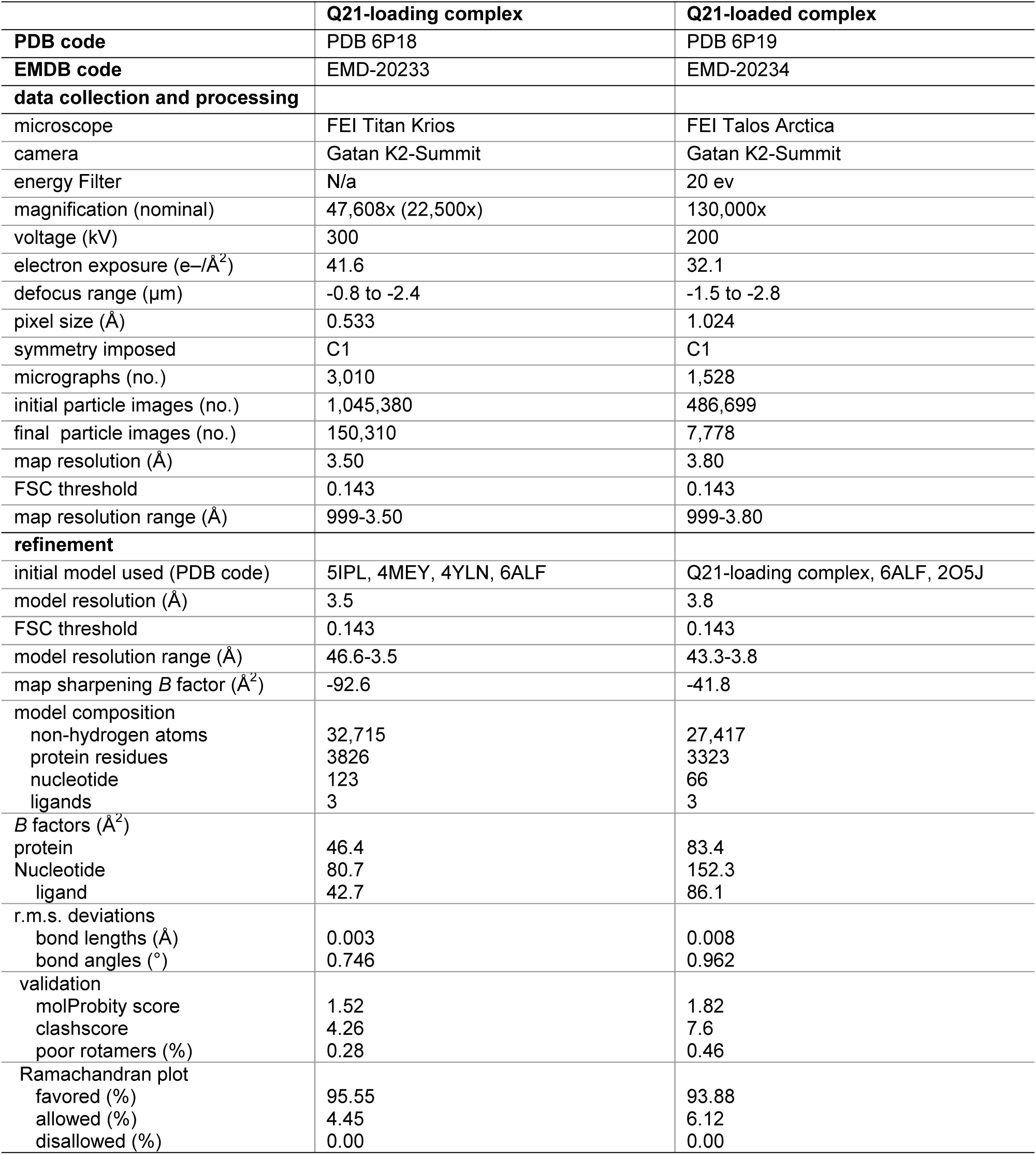
Data-collection, refinement, and validation statistics for cryo-EM structure of Q21-loading complex and Q21-loaded complex.

|  | Q21-loading complex | Q21-loaded complex |
| --- | --- | --- |
| **PDB code** | PDB 6P18 | PDB 6P19 |
| **EMDB code** | EMD-20233 | EMD-20234 |
| **data collection and processing** |  |  |
| microscope | FEI Titan Krios | FEI Talos Arctica |
| camera | Gatan K2-Summit | Gatan K2-Summit |
| energy Filter | N/a | 20 ev |
| magnification (nominal) | 47,608x (22,500x) | 130,000x |
| voltage (kV) | 300 | 200 |
| electron exposure (e-/Å<sup>2</sup>) | 41.6 | 32.1 |
| defocus range (µm) | -0.8 to -2.4 | -1.5 to -2.8 |
| pixel size (Å) | 0.533 | 1.024 |
| symmetry imposed | C1 | C1 |
| micrographs (no.) | 3,010 | 1,528 |
| initial particle images (no.) | 1,045,380 | 486,699 |
| final particle images (no.) | 150,310 | 7,778 |
| map resolution (Å) | 3.50 | 3.80 |
| FSC threshold | 0.143 | 0.143 |
| map resolution range (Å) | 999-3.50 | 999-3.80 |
| **refinement** |  |  |
| initial model used (PDB code) | 5IPL, 4MEY, 4YLN, 6ALF | Q21-loading complex, 6ALF, 2O5J |
| model resolution (Å) | 3.5 | 3.8 |
| FSC threshold | 0.143 | 0.143 |
| model resolution range (Å) | 46.6-3.5 | 43.3-3.8 |
| map sharpening *B* factor (Å<sup>2</sup>) | -92.6 | -41.8 |
| model composition |  |  |
| non-hydrogen atoms | 32,715 | 27,417 |
| protein residues | 3826 | 3323 |
| nucleotide | 123 | 66 |
| ligands | 3 | 3 |
| *B* factors (Å<sup>2</sup>) |  |  |
| protein | 46.4 | 83.4 |
| Nucleotide | 80.7 | 152.3 |
| ligand | 42.7 | 86.1 |
| r.m.s. deviations |  |  |
| bond lengths (Å) | 0.003 | 0.008 |
| bond angles (°) | 0.746 | 0.962 |
| validation |  |  |
| molProbity score | 1.52 | 1.82 |
| clashscore | 4.26 | 7.6 |
| poor rotamers (%) | 0.28 | 0.46 |
| Ramachandran plot |  |  |
| favored (%) | 95.55 | 93.88 |
| allowed (%) | 4.45 | 6.12 |
| disallowed (%) | 0.00 | 0.00 |

